# Label-free histological analysis of retrieved thrombi in acute ischemic stroke using optical diffraction tomography and deep learning

**DOI:** 10.1101/2023.02.22.529519

**Authors:** Yoonjae Chung, Geon Kim, Ah-Rim Moon, Donghun Ryu, Herve Hugonnet, Mahn Jae Lee, Dongseong Shin, Seung-Jae Lee, Eek-Sung Lee, Yongkeun Park

## Abstract

For patients with acute ischemic stroke, histological quantification of thrombus composition provides evidence for determining appropriate treatment. However, the traditional manual segmentation of stained thrombi is laborious and inconsistent. In this study, we propose a label-free method that combines optical diffraction tomography (ODT) and deep learning (DL) to automate the histological quantification process. The DL model classifies ODT image patches with 95% accuracy, and the collective prediction generates a whole-slide map of red blood cells and fibrin. The resulting whole-slide composition displays an average error of 1.1% and does not experience staining variability, facilitating faster analysis with reduced labor. The present approach will enable rapid and quantitative evaluation of blood clot composition, expediting the preclinical research and diagnosis of cardiovascular diseases.

## 1. Introduction

Acute ischemic stroke (AIS) is a fatal disease that requires immediate and appropriate treatment to prevent death or devastating sequelae. The composition of the thrombus, or blood clot, provides useful evidence in determining the appropriate treatment for AIS. The histological composition and structure of AIS thrombi offer insights into disease pathophysiology, such as pre-interventional migration, vascular origin, and thrombus age, and may be used to decide the best treatment options for patients. The thrombus composition is also related to the clinical outcomes of recanalization with thrombolysis or endovascular thrombectomy [1]. Studies have identified the pre-interventional migration, vascular origin, and age of thrombi from their histological composition and structure [2, 3], while others have differentiated cardioembolic from non-cardioembolic origin using the proportion of fibrin/platelets or red blood cells (RBCs) in thrombi [4, 5].

Conventional approaches to evaluating the histological composition of a thrombus require pathologists to manually screen stained thrombus sections under a bright-field (BF) microscope. Through microscopic examination, pathologists differentiate the type of thrombi into RBC-dominant or fibrin/platelet-dominant and report distinct features based on the stained thrombus slide [6-8]. However, color-based analysis of a thrombus slide is highly influenced by the staining quality and may lead to generalization problems due to staining variability [9]. Additionally, the fixation and staining procedures are labor-intensive and time-consuming, limiting efficiency and throughput.

In this study, we propose and experimentally demonstrate a label-free histological quantification framework to rapidly assess AIS thrombi using optical diffraction tomography (ODT) and a deep learning (DL) algorithm (Fig. 1). We leverage the label-free structural assessment of ODT with the statistical image recognition of deep learning to automatically characterize a whole unstained slide of the thrombus. ODT is a label-free 3D imaging method for cells and tissues that returns the refractive index (RI) distribution of the sample [10, 11]. The label-free nature of ODT simplifies sample preparation and enables efficient imaging, while also providing the ability to extract various quantitative biophysical properties from the RI distribution.

**Fig. 1.**
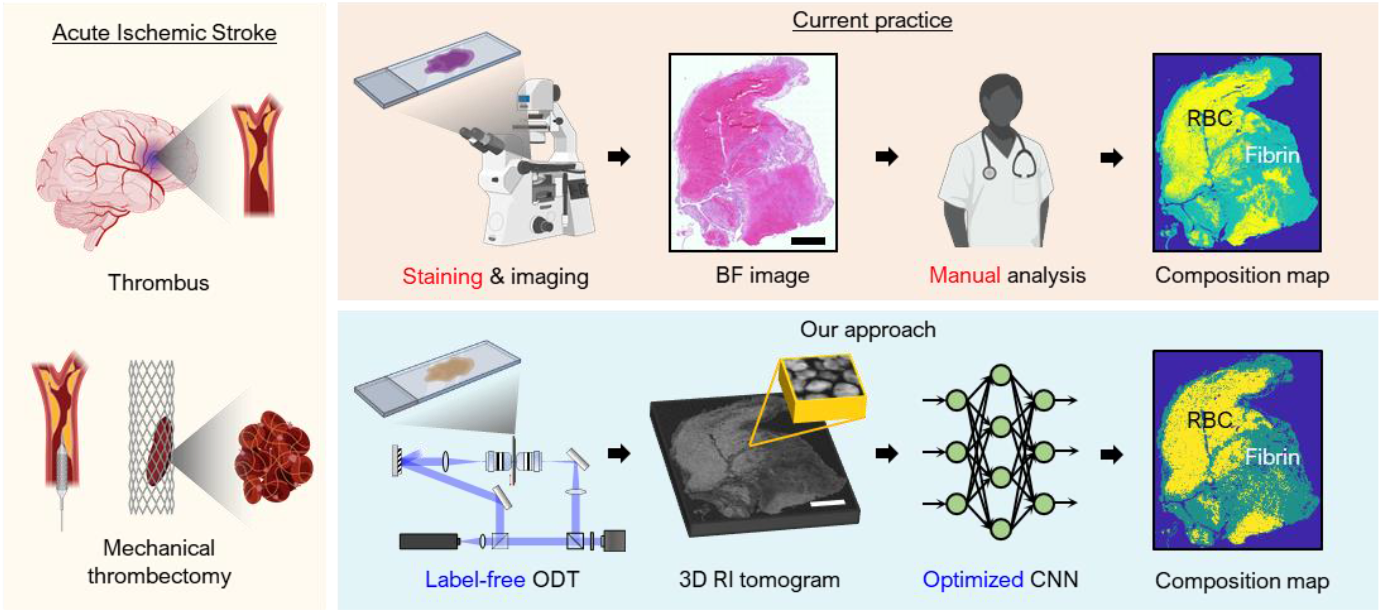
Proposed thrombus composition prediction compared to the current practice. Conventionally thrombus composition is manually analyzed though stained bright-field (BF) images. Our framework fusing ODT and DL provides label-free and fully-automated prediction of the composition. Scale bar = 500 μm.

This work exploits the high-resolution and quantitative 3D imaging capability of ODT, which extract various quantitative biophysical properties from the RI distribution [10]. These strengths of label-free imaging and ODT had been demonstrated in histological studies [12-17], including kidney cancer [18], kidney injury [19], and intestinal inflammation [20]. The use of refractive index as reproducible and quantitative imaging contrast has strengths when combined with DL. Recently, various cellular and subcellular RI distributions have been analyzed using the combination of HT and DL, including cellular segmentation [21-23], the detection of biological compartments [24-26] and domain translations [27, 28].

Our approach facilitates label-free and automated examination of an AIS thrombus by utilizing DL to predict the composition of the thrombus from a 3D RI tomogram. We validated the prediction from each of the three unstained thrombus slices by comparing to a successive slice that is Hematoxylin and Eosin (H&E) stained. An accuracy of 95% was achieved in the patch-wise classification when compared to board-certified pathologists’ annotation. Furthermore, the spatial distribution of thrombus composition was in high agreement with the annotation over the entire slide, highlighting the scope for whole-slide analyses. We expect that our study will enable rapid and quantitative evaluation of thrombus composition, aiding clinicians in treating AIS efficiently.

## 2. Methods

### 2.1 Sample preparation

Thrombi were retrieved from the middle cerebral artery by stent-based thrombectomy using a Solitaire stent (Covidien, Irvine, California) from a patient with AIS in Soonchunhyang University, Bucheon Hospital. The thrombi were sliced into sections (4-μm thickness) using a microtome. Two consecutive sections of a thrombus were visualized using two different imaging modalities: ODT and bright-field microscopy. Although the two consecutive sections may differ in microstructure, we assumed that they had nearly identical compositions of red blood cells (RBCs) and fibrin. One thrombus section was prepared with H&E staining, while the other was prepared without staining. Both thrombus sections were mounted on standard glass slides and sealed with coverslips (Fig. 2(a)).

**Fig. 2.**
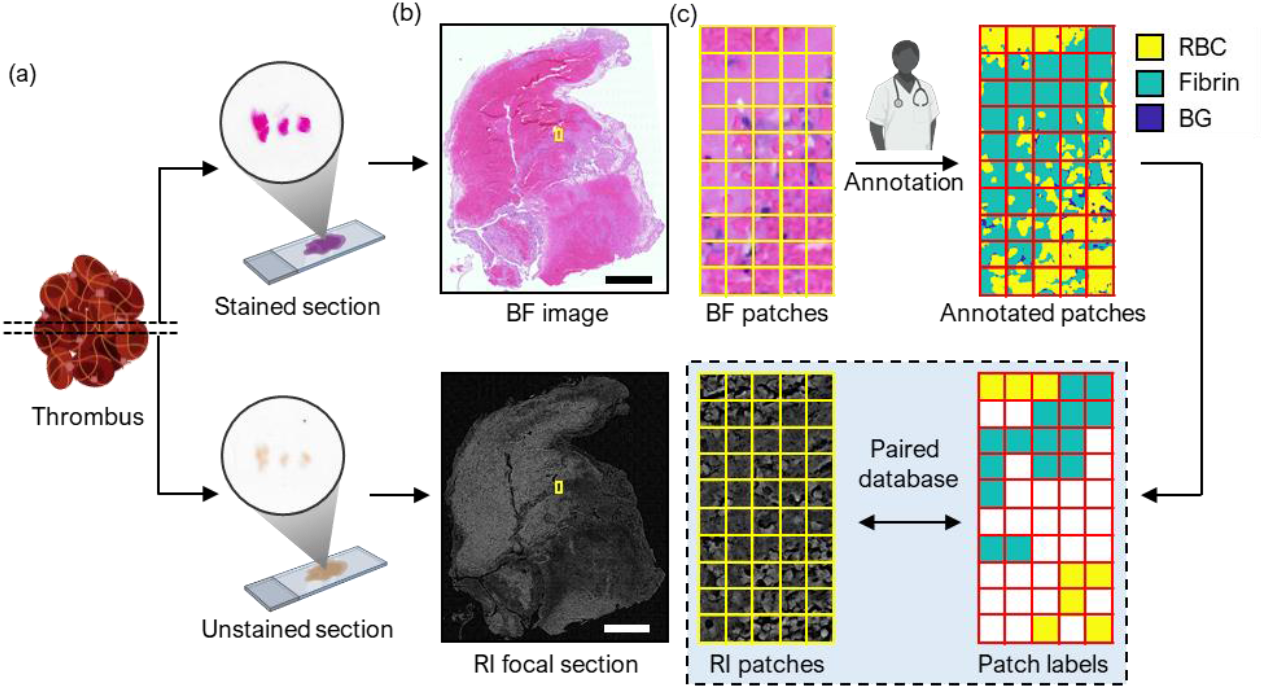
Flowchart of patch-wise annotation for RI images. (a) Two consecutive sections of a thrombus are retrieved, one of which is H&E stained. (b) The stained and unstained sections are respectively imaged with BF microscopy and ODT, and registered. A single focal section of the RI tomogram with the highest contrast is utilized. (c) The registered images are divided into patches. Each patch is labeled based on the annotation of a trained pathologist. Scale bar = 500 μm.

### 2.2 Optical measurement

The paired H&E stained and unstained slides of the thrombus were imaged with a BF slide scanner (Axio Scan.Z1; Zeiss; Jena, Germany) and ODT, respectively (Fig. 2(b)). An ODT system was based on Mach–Zehnder interferometry and equipped with a digital micromirror device (DMD) for high-speed angle scanning (Fig. S1). A blue continuous-wave laser (*λ*_0_ = 457 nm, Cobolt Twist, Cobolt) was split into a sample and reference beam using a beam splitter. The sample beam was modulated by the DMD for illumination angle control by impinging on a sample slide through a condenser objective (LUCPLFLN40X, NA = 0.6, Olympus). Then, the scattered fields from the sample at each illumination angle were collected by an objective lens (UPlanSAPO20X, NA = 0.75, Olympus) and interfered with the reference beam at the camera plane, generating a spatially modulated interferogram. Using field retrieval based on spectral filtering, complex field information consisting of amplitudes and phases was retrieved (Fig. S1). The 2D optical field information at various illumination angles was used to reconstruct a 3D refractive index (RI) tomogram based on Fourier diffraction theorem [29, 30]. Our setup had a lateral resolution of 0.17 μm and an axial resolution of 1.4 μm based on the Lauer criteria [31, 32].

To obtain whole-slide RI tomograms, multiple tomograms were stitched following the measurements in separate locations [33]. A phase-correlation algorithm [33, 34] was used to determine the relative position between overlapping tomogram tiles, and stitch them into tomograms. By automatically stitching the scanned tomograms, the ODT setup achieved a millimeter-scale FOV (10 mm^2^). The stitched tomogram had a size of approximately 10^4^ voxels in both lateral directions and 10^2^ voxels in the vertical direction. The BF images acquired using a slide scanner were 2D images. Hence, the 3D RI tomograms were converted to 2D by identifying the optimal focal plane. For training, RI values between 1.48 and 1.5 were normalized to a grayscale from 0 to 255.

We registered the RI and BF images to annotate each RI patch based on the corresponding BF patch. Subsequently, we adjusted the stained bright-field image to match the size and FOV of the unstained QPI. Rescale, rotation, skew, and distortions were applied to the bright-field images using Adobe Photoshop (Version 2019, Adobe Systems, San Jose, CA, USA).

### 2.3 Data annotation

Since the two axially consecutive images had nearly identical compositions of RBC and fibrin, we could label each RI patch (64 × 64 pixels) based on the H&E bright-field image, which had highly correlated information compared to the RI image. Experienced pathologists performed color-based semi-automatic segmentation on bright-field images using Adobe Photoshop (Version 2019, Adobe systems, San Jose, CA, USA), used as a standard reference technique. H&E-stained bright-field images were annotated by two board-certified pathologists to obtain ground truth information. Each pixel of the registered bright-field images was annotated as one of three types: an RBC-rich area, fibrin/platelet-rich area, or background. The ground-truth histological composition of each thrombus sample was determined by counting the number of RBC and fibrin pixels from the pathologist-annotation result.

After annotating the H&E-stained bright-field image, we divided the unstained QPI and bright-field images into small patches of 64 × 64 pixels (FOV 10.88 μm × 10.88 μm). For each QPI image patch, there was a corresponding bright-field image patch at the same position that had already been annotated. To predict the histological composition with DL, we train a DL model to classify small RI tomogram section patches into three categories: RBC, fibrin, and background. For the training dataset, we prepared 64 ×64 pixels QPI image patches, categorized into three components (RBC, fibrin, and background) using the information from the registered and annotated BF image patches (Fig. 2). We designed a label generation and patch selection rule to assign the class of the image patch (0: RBC, 1: fibrin, and 2: background) and exclude ambiguous patches that contain composite components (Fig. 2(c)). For every patch, the initial class was determined by counting the number of classified pixels in each class (RBC, fibrin, and background) and assigning the class indicated by the most pixels.

To provide a suitable database for DL, we selected cases where over 80% of the pixels in the patch indicated a certain class to form an accurate training set. Patches that contain many pixels that do not belong to the assigned class are undesirable for DL training. For example, a patch that contained 40% of RBC pixels, 30% of fibrin pixels, and 30% of background pixels would be excluded from the training set. Note that the selected patch size, 10.88×10.88 μm^2^, was sufficiently small to prevent the exclusion of too many patches that did not have dominating components while selecting 76% of the total patches for the training set.

### 2.4 Deep learning and optimization

The proposed deep neural network consists of convolutional layers, pooling layers, ReLU activation layers, and fully connected layers. It takes a 64 × 64 RI image patch and classifies it into one of three subtypes (RBC, fibrin, and background). The network extracts various spatial features of each patch using contracting convolutional operations (7 × 7, 3 × 3) with nonlinear operations. The final label is determined using a fully connected layer. Batch normalization is applied at every stage before the ReLU activation to accelerate training speed and improve regularization.

The labeled image patches were divided into training, validation, and test sets at a ratio of 7:1.5:1.5. Data augmentation techniques, including random rotation and flips, were used only during the training stage. We set the cross-entropy loss based on the SoftMax function. For parameter learning with backpropagation, we used the Adam optimizer with β_1_ = 0.9, β_2_ = 0.999, a learning rate of 1×10^−5^, and a batch size of 256 [35]. We used graphics processing units (GPU) (TESLA P40, NVIDIA) with CUDA Toolkit 10.0 (NVIDIA) for training. After 200 epochs of training, we selected the optimal model by early stopping, evaluating the validation loss for every epoch. We used the Python 3.7.2 environment with PyTorch version 1.8.1.

### 2.5 Inference of histological composition

To test our trained network, we imaged the RI of an unseen thrombus and quantified its composition. The trained DL model classified each patch (64 × 64) of the RI image as RBC-rich, fibrin-rich, or background. For rapid inference, parallel input with a batch size of 1,024 was used. Based on the classification results, the thrombus composition was calculated by specifying the proportions of RBC and fibrin. The spatial distribution of the RBC and fibrin components could also be visualized by coloring the RI patch position with pseudocolors according to the classification result and arranging the patch to the original spatial position.

Using patch-by-patch inference makes this method robust to local mismatches during image registration. The DL model classifies the patch by evaluating the overall features and is tolerant to a small portion of mislabeled pixels.

## 3. Results

### 3.1 Microscopic measurements of the thrombus

We first measured the BF image of a H&E-stained thrombus slide and the RI distributions of an unlabeled consecutive slice using a BF slide scanner and an ODT system, respectively (Fig. 3; see Methods for details). The BF image revealed a clear distinction between RBC and fibrin (Fig. 3a). The RI distribution measured using the ODT provided high-resolution subcellular features and exhibited characteristic RI values for the RBC and fibrin area.

**Fig. 3.**
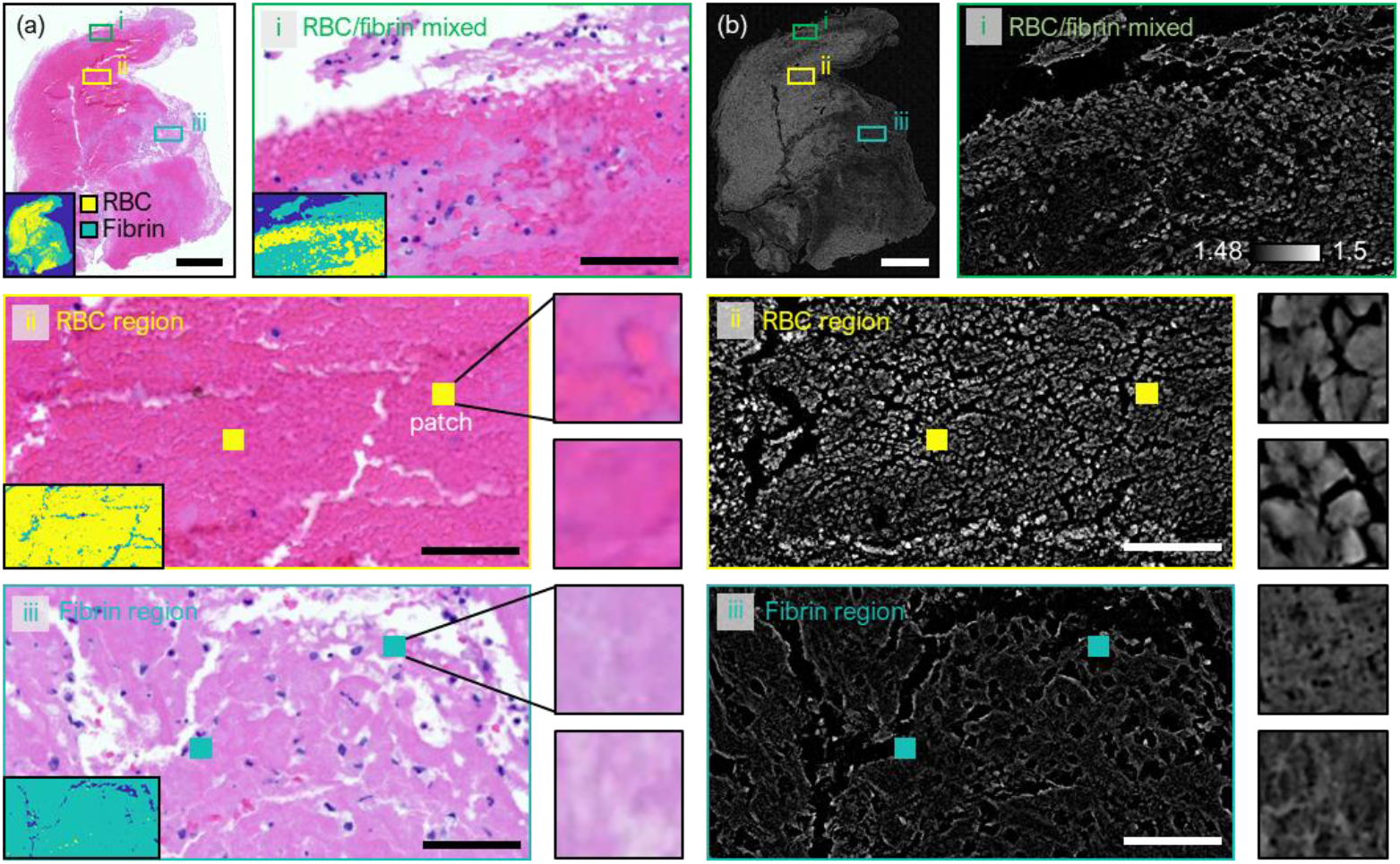
Optical measurement. (a) BF image and (b) RI focal section. The boxed regions i, ii and iii denote mixed, RBC, and fibrin regions respectively. The size of each patch is 10.88 μm × 10.88 μm. Scale bars = 500 μm (boxed regions) and 50 μm (patches).

To validate the quality of the RI reconstruction, we visually compared the thrombus structure in the paired RI tomogram section and BF image. Figure 3 shows the BF and RI images of thrombus sample 3 at various magnifications: whole-slide level, multiple patches level, and individual patch level. The RI tomograms of the thrombus show various RI value distributions ranging from 1.48 to 1.5. Compared to the BF image, the RI tomogram section visualizes RBCs and fibrin structures of thrombus components better. The contrast of RI is directly related to the concentration of material [36, 37]. In Figure 3, the magnified region of interest (ROI) of sample 3 is presented. In the RI tomogram section (Fig. 3(b)), some borders of individual RBC are clearly shown even in tightly compressed region, whereas in the corresponding BF image (Fig. 3(a)), the borders of individual RBCs are unclear. Some RBC borders in the RI image possibly indicate polyhedrocytes, which are tightly packed red blood cells (RBCs) with polyhedral shapes formed when blood clots contract. In the fibrin region, (iii) of Fig. 3, the pores in the meshwork of fibrin fibers are also better observed in the RI tomogram. By contrast, the boundary of fibrin pores cannot be identified in the BF image.

### 3.2 Patch-wise composition prediction based on DL

Our dataset for DL-based prediction consisted of 166,887 patches of focal RI sections, with a FOV of each patch at 10.88 μm × 10.88 μm. For each slide, the patches were randomly separated into the training, validation, and test sets in a ratio of 7:1.5:1.5.

The DL classifier was trained using the training and validation datasets and blind-tested using the test dataset (Fig. 4(a)). The optimal model was chosen based on the validation loss during DL training, and the learning curve is presented in Fig. 4(b). The resulting test accuracy of the DL model was 95.1% (Fig. 4(d)). A major error in the patch-wise classification was the misclassification of RBC into fibrin, accounting for 10.4% of all the RBC patches in the test set. This error could be due to the limited correspondence between the two consecutive thrombus sections, resulting in imperfectly mixed regions in the registered patch annotation. Additionally, the rate of confusing fibrin with the background was higher (2.97%) than that of confusing RBC with background (0.577%). This difference in background-related error is consistent with our observations of thin fibrin structures at the edge of the slide and the relatively low RI distribution of fibrin.

**Fig. 4.**
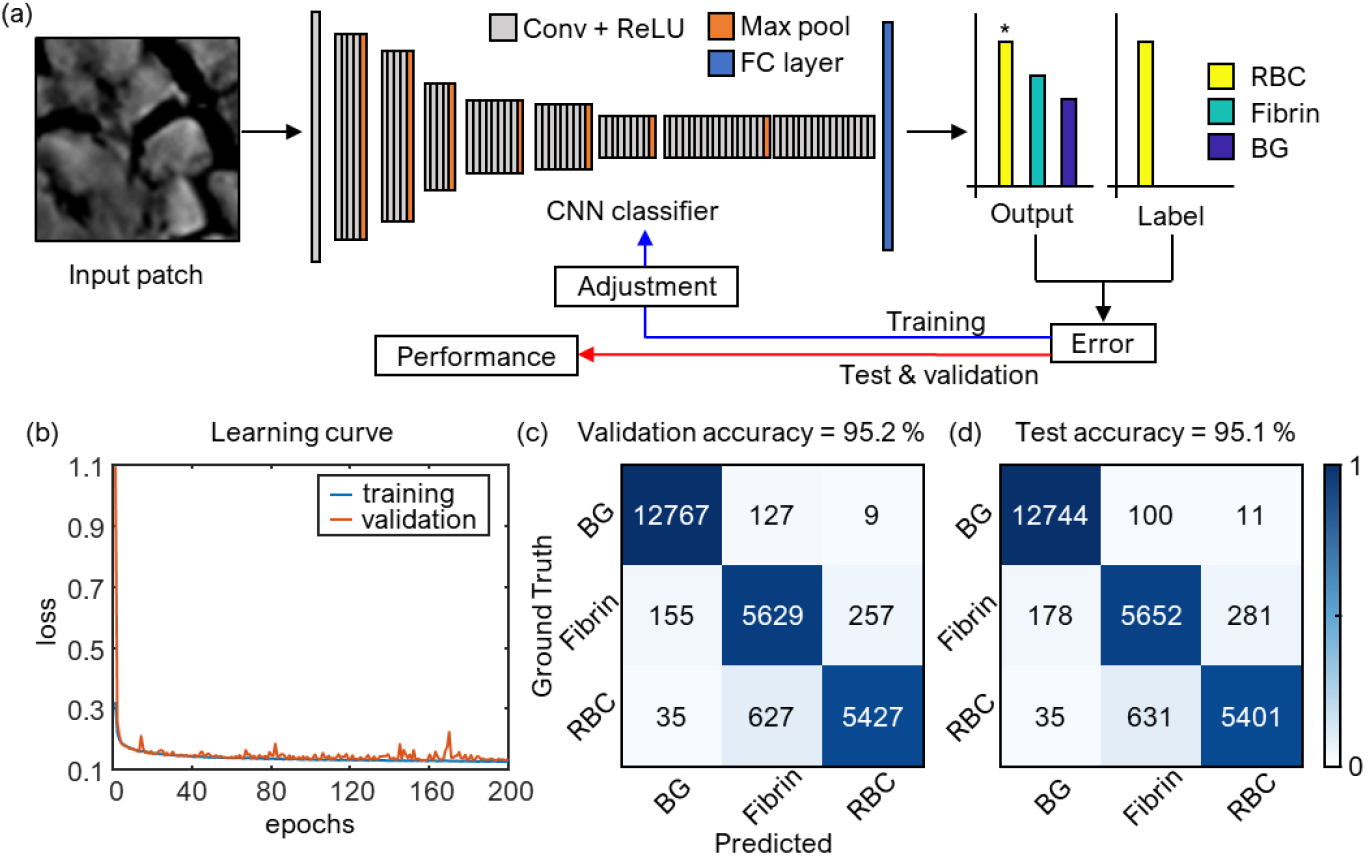
Deep learning classification of RI patches. (a) The architecture of the CNN classifier for RI patches. (b) The loss curve for the training set and the validation sets during 200 epochs of training. (c)-(d) validation and test confusion matrices.

### 3.3 Whole-slide analysis of thrombus composition

To further assess the performance of the method in analyzing whole-slide images, the class of each patch was identified and stitched together to create a large field-of-view thrombus RI image. The patch classification results were used to create a virtually colored patch image and an unstained image, which were compared to the ground truth annotation map based on H&E-stained images. Figure 5 shows the results for four different regions, including slides 1, 2, and 3, and a region of interest (ROI) from slide 3. Despite the differences in resolution due to the patch-wise processing, the prediction based on DL and ODT showed a similar spatial distribution of RBC and fibrin regions as the ground truth. The sizes of the regions varied from 101 × 334 μm^2^ for the ROI to 2.71 × 5.58 mm^2^ for slide 1.

**Fig. 5.**
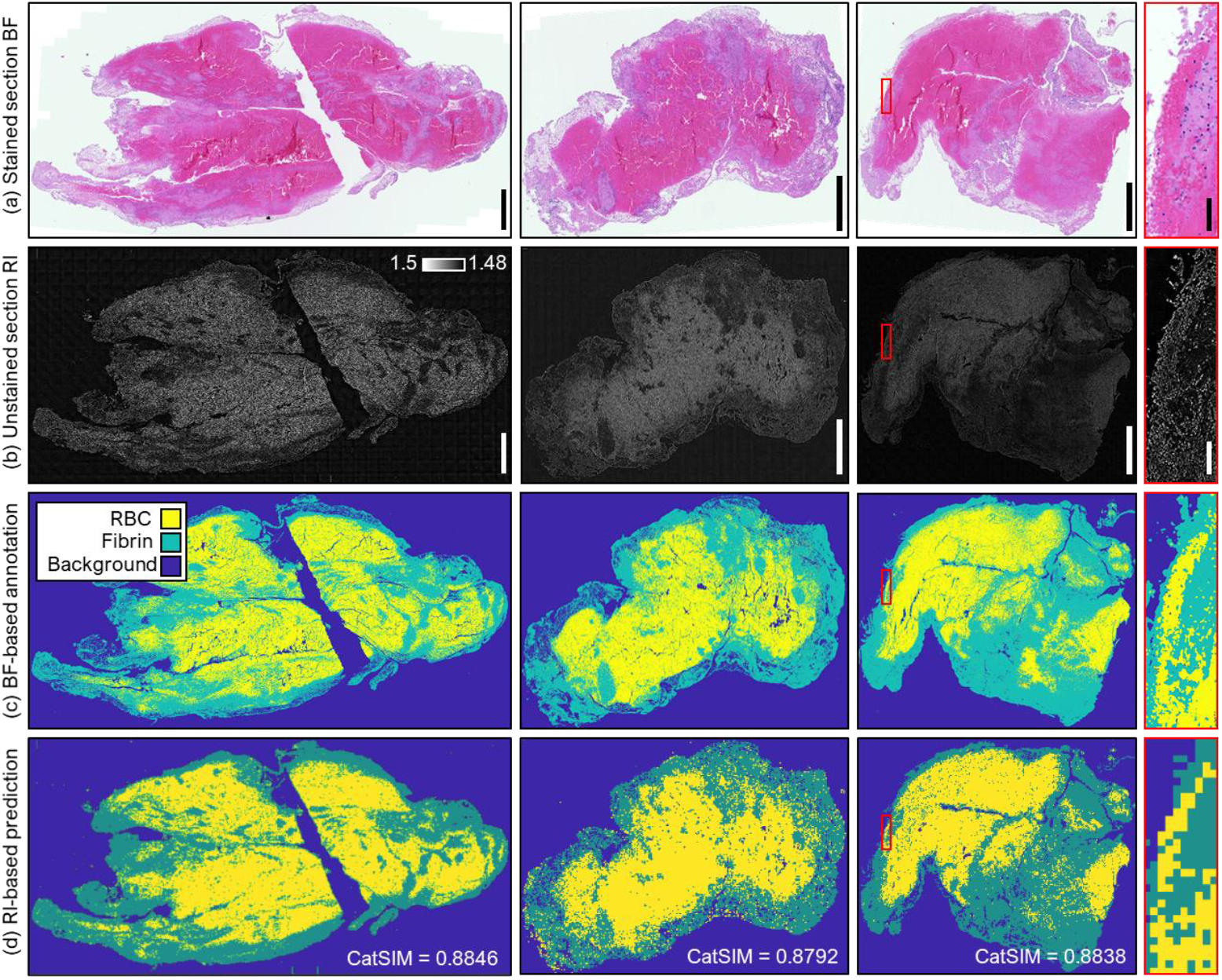
Whole-slide analysis using the proposed method. (a) H&E-stained slide images, (b) corresponding RI sections, (c) annotation by pathologists from (a), and (d) CNN-based prediction from (b). The first three rows illustrate three different slides, and the last row is a magnified view of the third slide. Scale bars = 500 μm (first three columms) and 50 μm (last column).

To quantitatively evaluate the similarity between the predicted and ground truth images, the categorical image similarity metric (CatSIM) was computed [38]. CatSIM measures the similarity in RBC and fibrin distribution and uses the vector of proportions of each class and categorical variance, which is related to the diversity index. The terms for categorical luminance, contrast, and structural similarity were calculated to determine the CatSIM value (Supplement 1). The CatSIM values for all three samples were close to 0.9, indicating that the DL prediction successfully inferred the thrombus structure and accurately identified the RBC-rich and fibrin-rich areas.

### 3.4 Thrombus composition ratio and cross-validation

The DL model predicted the quantitative composition of the patient’s thrombus sample slides at the whole-slide level (Fig. 6(a)). The average error between the predicted composition and the ground truth was 1.1%, with a maximum error of 2.6%. We employed a slide-level cross-validation scheme to evaluate the DL model without bias in a limited number of samples. Each whole-slide sample provided at least ∼3×10^4^ patches, which is sufficient for DL training. There was slight variability in the prediction results based on different training data (Fig. 6(b)). CNN2, which was trained with slide 2, tended to predict a higher ratio of fibrin than the ground truth (1.0% and 2.5%). The slide that was the least accurate was slide 1. Although the error was maintained under a certain level, the cross-slide error is thought to be largely a result of the difference between the adjacent slides or the variable nature of staining.

**Fig. 6.**
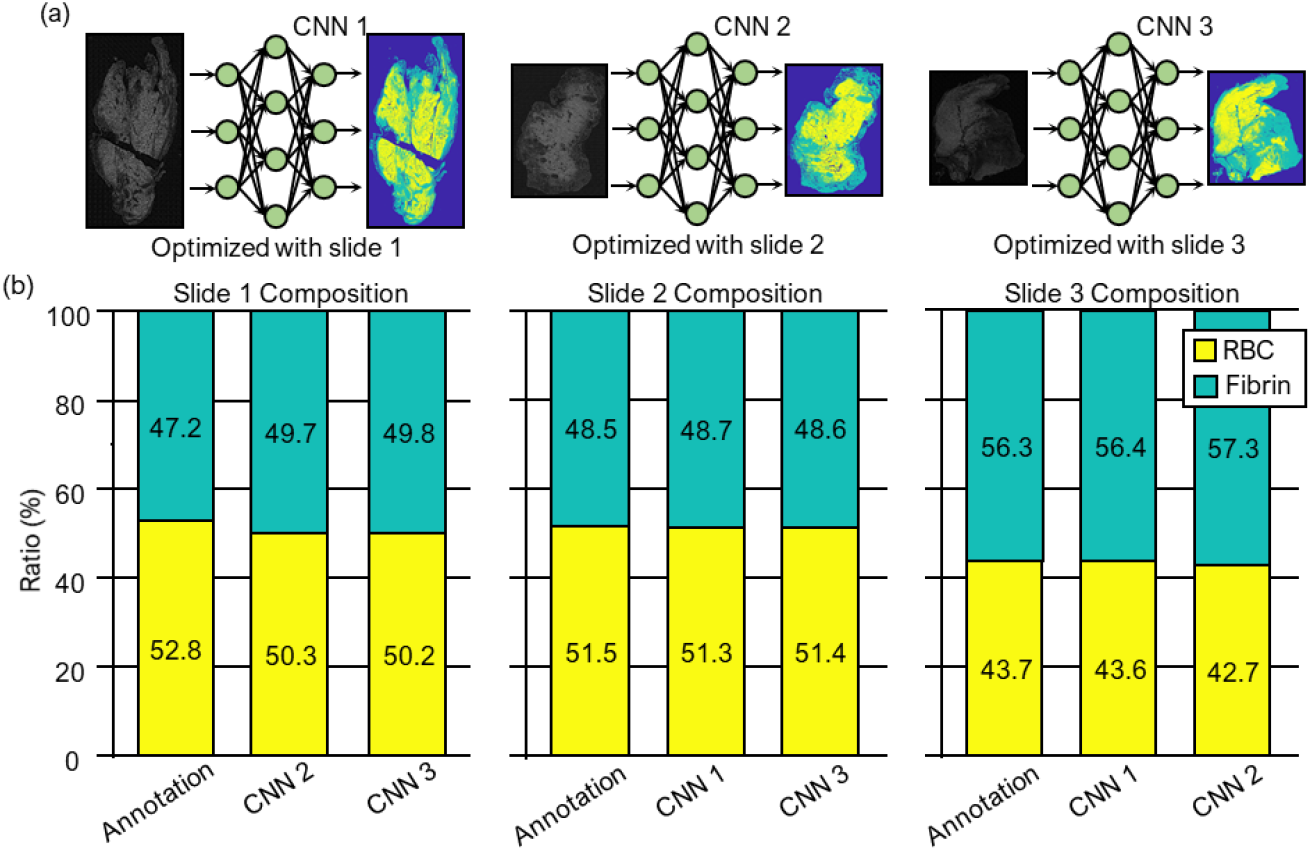
Variation of the prediction based on the training data. (a) CNN classifiers optimized based on different slides. (b) thrombus composition results corresponding to different CNN classifiers.

## 4. Discussion

This study presents a DL-based approach to classify the histological composition of thrombi using label-free ODT images. The DL model achieved over 95% accuracy in classifying patches into RBC, fibrin, and background, enabling the prediction of thrombus composition at whole-slide level. The use of ODT provided higher resolution and contrast compared to conventional BF microscopy followed by H&E staining. This label-free approach also allowed for rapid histological analysis without staining and reduced the variability of staining-dependent color distribution. The present work demonstrates the rapid label-free detection a spatial distribution map of thrombus composition that can be used to assess the thrombus response to thrombectomy procedure [39].

The results shown here can also be utilized for in-depth investigation of a thrombus slide. The RI contrast and subcellular spatial resolution provided by ODT facilitate a more detailed assessment of thrombus structures compared to BF microscopy followed by H&E staining. Pores within the fibrin network were more clearly observed in ODT than in BF images. The border and texture of individual RBCs also appeared sharper in ODT. Additionally, ODT distinguished structures that were not differentiated by BF microscopy, such as regions of compressed RBCs tentatively identified as polyhedrocytes. Polyhedrocytes may lead to more thrombolysis-resistant thrombi by forming an impermeable layer that blocks fibrinolytic enzymes from diffusing inside the thrombi. These observations indicate that ODT measurement could potentially identify indicators of thromboembolism [40].

Additionally, accounting for volumetric information could provide a more accurate determination of thrombus composition due to intra-thrombus heterogeneity. Ground truth generation could also be improved by utilizing immunofluorescence or immunohistochemistry. The DL design could be modified to carry out segmentation instead of patch classification for better reflecting the high-resolution geometry provided by ODT. Using identical slides for ODT and bright-field imaging may also improve pixel-level registration and provide more accurate supervised learning for segmentation networks.

The proposed framework has the potential to complement the current clinical routine by providing rapid on-site evaluation (ROSE) of thrombi. H&E staining and inspection of thrombi remains a solid gold standard in most clinical institutes owing to the effectiveness that results from abundant domain knowledge. Our noninvasive test can be integrated into this routine slide inspection workflow without perturbing or prohibiting the existing processes. The use of label-free ODT enabled us to reduce the time and associated effort of staining. Our approach does not require any staining and can directly assess the unlabeled specimen, allowing rapid histological analysis of patients with AIS. Our approach can also provide consistent results compared with conventional methods that suffer from staining variability and interpreter fatigue. The conventional composition assessment by practitioners depends heavily on the color distribution which results from staining [7]. As the staining procedures rely on human or environmental conditions, the color distribution may be significantly different [9, 41, 42]. By contrast, it does not apply to our label-free approach that measures RI, an intrinsic physical quantity of the native sample.

There are technical limitations in this work, such as the bulkiness and complexity of the ODT hardware implementation, the limit in imaging speed arising from the small FOV compared to a whole slide, and the acquisition rate of the image sensor. However, recent developments in non-interferometric ODT using low-coherence sources and engineering approaches such as multiplexing can mitigate these issues [43, 44]. Other possible engineering approaches include multiplexing which expands the measurement bandwidth by exploiting the polarization [45] or spectral dimension [46].

## 5. Conclusions

We proposed and experimentally demonstrated a rapid and fully automated prediction of thrombus composition using label-free refractive index (RI) tomography and deep learning. Our proposed convolutional neural network model accurately classified thrombus subtypes for each label-free RI patch with high accuracy (>95%), affording quantitative analysis of unstained thrombus slides. We believe that this approach, which does not require any staining or manual inspection, will significantly accelerate thrombus biopsy procedures for diagnosing ischemic stroke and related diseases.

In future work, we may extend our framework to fully exploit the 3D information of the RI tomogram, which could enhance the accuracy of the thrombus composition inference. Our DL-based approach using label-free ODT images presents a promising alternative to conventional staining-based methods for histological analysis of thrombi, which can also be further extended to identify and classify cell types in tissue slides. The proposed framework has the potential to provide more efficient and consistent analysis while complementing the existing clinical routine.

## Acknowledgments

We thank the patients and staffs of Soonchunhyang University Hospital Bucheon for providing the AIS thrombus slides.

## Data availability

Data underlying the results presented in this paper are not publicly available at this time but may be obtained from the authors upon reasonable request.

## Supplemental document

See Supplement 1 for supporting content.

